# Glycogen Synthase Kinase-3β Regulates Cellular Prion Protein Levels

**DOI:** 10.64898/2026.08.21.746199

**Authors:** Kathryn S. Beauchemin, Anna M. Schmoker, Joel C. Watts, Surachai Supattapone

## Abstract

The normal cellular prion protein (PrP^C^) is an essential substrate in all forms of prion diseases and a receptor for Aβ oligomers in Alzheimer’s disease. However, it is not fully understood how cells regulate PrP^C^ levels. Recently, we identified glycogen synthase kinase-3β (GSK-3β) as a potential regulator of PrP^C^ levels in a whole genome knockout screen. Here, we show that both cell surface and total PrP^C^ levels can be reduced either by siRNA-mediated *Gsk3b* (but not *Gsk3a*) knockdown or by CRISPR-mediated *Gs3b* knockout. Whole cell mass spectrometric analysis showed that PrP^C^ was the 60^th^ most significantly reduced protein (out of 7227 total proteins detected) in *Gsk3b* knockout cells, compared to wild-type cells. Two different GSK-3 inhibitors, laduviglusib (CHIR-99021) and AZD-1080, reduced PrP^C^ levels in mouse CAD5 and human BE(2)-C cells, both in undifferentiated and differentiated states. PrP^C^ levels were similarly reduced by cycloheximide treatment in both *Gsk3b* knockout and WT cells, indicating that GSK-3β regulates PrP^C^ levels through a post-translational mechanism. Finally, treatment with either laduviglusib or AZD-1080 reduced PrP^Sc^ levels in CAD5 cells infected with three different rodent prion strains. Overall, the results reveal that GSK-3β activity controls PrP^C^ levels in living cells, revealing a novel regulatory mechanism and promising therapeutic target.

## Introduction

Prion diseases and Alzheimer’s Disease (AD) are fatal neurodegenerative disorders characterized by the misfolding and aggregation of specific proteins within the brain. In all forms of prion disease (inherited, sporadic, and infectious), the cellular prion protein (PrP^C^) is essential for prion propagation, serving as a substrate for conversion into all pathogenic conformers, collectively termed PrP^Sc^[1]. There is also evidence that PrP^C^ plays a key role in AD as a high-affinity receptor for neurotoxic amyloid-β (Aβ) oligomers, mediating synaptic dysfunction and memory impairment[2]. Reducing PrP^C^ expression or availability has been shown to improve survival in animal models of both prion disease and AD[3, 4]. This dual role in both prion and Alzheimer’s pathogenesis positions PrP^C^ as a uniquely compelling therapeutic target for modifying disease progression across these otherwise distinct disorders.

Despite this strong rationale, current approaches to lower PrP^C^ levels face practical limitations. Genetic knockdown strategies (e.g., antisense oligonucleotides or RNAi) are often hampered by limited CNS penetration (particularly to deep-lying brain regions such as the thalamus and hippocampus), off-target effects, and the need for invasive administration routes[5, 6]. Biological therapeutics, such as antibodies, similarly struggle with blood-brain barrier (BBB) permeability and often require repeated parenteral dosing. Therefore, it would be desirable to be able to reduce PrP^C^ levels by using small molecule compounds that can penetrate the BBB. An obstacle to developing such compounds is that the signaling pathways responsible for regulating cellular PrP^C^ levels remain poorly understood.

We recently conducted a genome-wide CRISPR knockout (KO) screen to identify regulators of cell surface PrP^C^ expression in CAD5 cells (a differentiable catecholaminergic murine CNS cell line highly susceptible to multiple prion strains[7]), measured by fluorescence-activated cell sorting (FACS)[8]. Intriguingly, this screen identified *Gsk3b* (but not its closely related paralog *Gsk3a*) as a potential positive regulator of PrP^C^ levels. *Gsk3b* encodes glycogen synthase kinase-3β (GSK-3β), an enzyme that phosphorylates tau and has been implicated in the pathogenesis of neurodegenerative diseases, particularly AD[9]. Since GSK-3β can be targeted with brain-penetrant small molecules[10–12], we sought to validate and characterize its ability to modulate cellular PrP^C^ levels.

## Materials and Methods

### Cell lines and cell culture

WT CAD5 cells were kindly provided by Charles Weissmann (Scripps Florida, Jupiter, FL, USA). CAD5 cells genetically engineered using CRISPR/Cas9 to lack endogenous mouse PrP expression and engineered to stably express either hamster PrP (CAD5-PrP^-/-^(Ha)) or bank vole M109 PrP (CAD5-PrP^-/-^(BV^M^)) were kindly provided by Joel Watts (University of Toronto, Toronto, Canada)[13, 14].

CAD5 cells were maintained in Opti-MEM I Reduced-Serum Media with GlutaMAX (Gibco, Waltham, MA, USA) supplemented with 10% HyClone Bovine Growth Serum (BGS) (Cytiva, Marlborough, MA, USA) and 1X penicillin/streptomycin (Corning, Corning, NY, USA). Prion-infected CAD5 cells were cultured using 0.2X penicillin/streptomycin. To differentiate CAD5 cells, cells were rinsed with PBS and complete media was replaced with DMEM:F12 (Gibco) supplemented with 50 ng/mL sodium selenite (Sigma-Aldrich, St. Louis, MO, USA). Differentiation was considered complete after 4 days incubation in the differentiation media.

BE(2)-C human neuroblastoma cells (ATCC CRL-2268) were cultured in a 1:1 mix of EMEM (ATCC) and F-12 (Gibco) media supplemented with 10% fetal bovine serum (Cytiva) at 37°C with 5% CO_2_ until >90% confluent then split 1:4-1:10 with 0.05% Trypsin / 0.53 mM EDTA (Corning). BE(2)-C differentiation was performed by adding all-trans retinoic acid (Thermo Fisher Scientific, Waltham, MA, USA) to a final concentration of 10 μM in complete growth media and was considered complete after 4 days of incubation.

Cell lines were routinely monitored for mycoplasma contamination using the LookOut Mycoplasma PCR Detection Kit (Sigma-Aldrich, St. Louis, MO, USA).

### GSK-3 inhibitor treatments

Unless otherwise noted, all experiments involving cell treatment with GSK-3 inhibitors underwent treatment at the indicated final concentration in complete cell culture media, with media/drug refreshes every 24 hr.

Laduviglusib (MedChemExpress, HY-10182) and AZD1080 (MedChemExpress, HY-13862) were prepared as 20 mM stock solutions in DMSO (Sigma-Aldrich, D2650). Both drugs were aliquoted and frozen at manufacturer’s recommended storage temperature. Thawed aliquots were discarded after use to avoid repeated freeze/thaw.

### Flow cytometry for surface PrP^C^

All flow cytometry experiments were performed on a Northern Lights flow cytometer (Cytek, Fremont, CA, USA).

To detect surface PrP^C^ in CAD5 cells, the anti-PrP antibody 6D11 (Biolegend, San Diego, CA, USA) was used at 0.8 µg per sample. The anti-PrP antibody 4D5:PE (Invitrogen, 12-9230-42) was used at a 0.125 µg per sample to detect surface PrP^C^ in BE(2)-C cells. Each sample contained approximately 1 million cells in 100 µL PBS/2% FBS (staining buffer). Primary incubations were performed at 4 °C for 1 hr, followed by three washes in PBS, with final resuspension to 100 µL in staining buffer for secondary staining or in 500 µL staining buffer for flow cytometry.

Anti-Mouse IgG2a Secondary Antibody:PE (Invitrogen, 12-4210-82) was used as the secondary antibody at the manufacturer’s recommended dilution for CAD5 staining. Secondary incubations were performed at 4 °C for 30 min, followed by three washes in PBS, with final resuspension in 500 μL staining buffer.

FlowJo software (BD Biosciences, Franklin Lakes, NJ, USA) was used for all flow cytometry analysis. Sequential gating using FSC and SSC characteristics were performed to identify single cell populations. Hoechst 33342 (ThermoFisher Scientific) was used at 5 μg/mL, 37 °C for 45 min, followed by two washes in PBS, to identify differentiated BE(2)-C cells containing DNA. Median fluorescence intensities of the resulting single cell populations were used for comparisons between populations and treatments.

### Prion infection of cultured cells

WT CAD5 cells were infected with mouse prion strains 22L and RML via incubation with brain homogenate from terminally ill animals. 10% (w/v) brain homogenate was prepared by weighing brains of terminally ill mice infected with either the 22L or RML strains and homogenizing for 30 sec in sterile 1X PBS (Corning) using a 4-Place Mini Bead Mill Homogenizer (VWR, Radnor, PA, USA) and sterile 1.4 mm ceramic beads (VWR,10158-552). 10% brain homogenate was stored at -80°C until use. WT CAD5 cells were seeded at a density of 1 x 10^5^ cells per well in a 12-well tissue culture plate (Corning) and incubated at 37°C, 5% CO_2_ overnight. 10% brain homogenate was thawed on ice, centrifuged at 400 x *g* for 30 sec at 4 °C twice to pellet debris, then diluted to 0.5% (w/v) brain homogenate in complete cell culture medium. Cell culture medium was replaced with the medium containing 0.5% (w/v) brain homogenate. Cells were exposed to the inoculum for 72 hr, then passaged at a 1:4 or 1:5 dilution every 2-3 days. Cells were passaged for one month prior to expansion, freezing, and analysis of prion infection status.

CAD5-PrP^-/-^(Ha) cells stably infected with Hyper were generated as described by Bourkas et al.[13].

### Cell lysis and immunoblotting

To immunoblot for PrP^C^, cells were washed 2X with PBS, lysed in lysis buffer (50 mM Tris, pH 8.0, 150 mM NaCl, 0.5% (w/v) sodium deoxycholate, and 0.5% (v/v) Nonidet P-40), collected, and centrifuged at 250 x *g* for 30 sec to pellet DNA. Lysate supernatant was transferred to fresh tubes. Protein concentrations were determined using a Pierce BCA Protein Assay Kit (Thermo Fisher Scientific) and 30-50 µg total protein was loaded per lane.

To check prion infection status, confluent cells were lysed as described above. After BCA analysis, lysates were digested with 20 µg/mL Proteinase K (PK) (Roche, Basel, Switzerland) for 1 h at 37 °C with 350 rpm shaking. Digestions were stopped by the addition of phenylmethylsulfonyl fluoride (PMSF) (Sigma-Aldrich) to a final concentration of 2 mM. Samples were then ultracentrifuged at 100,000 x *g* for 1 hr at 4 °C in a Sorvall Discovery M120 SE Micro-Ultracentrifuge with an S45-A rotor (Thermo Fisher Scientific) in safe-lock tubes (Eppendorf, Hamburg, Germany). Pellets resulting from either method were resuspended in 60 µL Milli-Q water plus 20 μL of 6X Laemmli SDS sample buffer (Bioland Scientific LLC, Paramount, CA, USA) containing 9% (v/v) beta-mercaptoethanol (BME) (Gibco) and boiled for 15 min at 95 °C. All PrP^Sc^ samples were normalized such that the same amount of total protein (500-1000 µg total protein pre-digest) was loaded between samples on each blot.

SDS-polyacrylamide gel electrophoresis (PAGE) was performed using 1.5 mm 12% polyacrylamide gels with an acrylamide/bisacrylamide ratio of 29:1. The gel was transferred to a methanol-charged polyvinylidene difluoride membrane (Millipore Sigma, Burlington, MA, USA) using a Transblot SD semidry transfer cell (Bio-Rad Laboratories, Hercules, CA, USA). The transfer was set at 2.5 mA/cm^2^ for 45 min. To visualize PrP signal, the membrane was blocked in 5% (w/v) nonfat dry milk (Nestlé, Vevey, Switzerland) in TBST (10 mM Tris, pH 7.1, 150 mM NaCl, 0.1% Tween 20) for 1 hr at 4 °C. The blocked membrane was then incubated overnight at 4 °C with anti-PrP 6D11 primary antibody, washed three times for 10 min in TBST, then incubated for 1 h with horseradish peroxidase-labeled sheep-anti-mouse secondary antibody (Cytiva). Membrane was then washed four additional times for 10 min each in TBST. For additional labeling, membranes were stripped for 15 min at 37 °C, rinsed under running dH_2_0, blocked in 5% (w/v) nonfat dry milk, and labeled with anti-beta catenin antibody CST9562 and/or anti-beta Tubulin antibody CST15115 and horseradish peroxidase-labeled goat-anti-rabbit secondary antibody (BioRad Cat # 1706515). Blots were developed with SuperSignal West Femto (Thermo Fisher Scientific) chemiluminescence substrate, and images were captured digitally using an Azure 600 (Azure Biosystems, Dublin, CA, USA) imaging system. Relative molecular masses were determined by comparison to PageRuler Plus Prestained Protein Ladder (Thermo Fisher Scientific).

### siRNA knockdown

All siRNA experiments utilized *Silencer* Select siRNAs (ThermoFisher) and were performed using undifferentiated CAD5s in 6-well tissue culture plates (Corning). Cells were seeded to achieve 60% confluence for transfection and were allowed to adhere for a minimum of 24 hr before transfection. All knockdowns were performed using a combination of two individual targeting siRNAs (ThermoFisher, *Gsk3a*: Cat # s121110, s1121111, *Gsk3b*: Cat # s80826, s80827). Nontargeting (scrambled) control siRNA (ThermoFisher Cat # 4390843) was used at the same final molar concentration as the combined targeting guides. For each well, 9 μL of Lipofectamine RNAiMAX transfection reagent (ThermoFisher) was diluted in 150 μL Opti-MEM. In a separate tube, 1.5 μL of each 10 μM siRNA stock (15 pmol per siRNA, 30 pmol total) was added to 150 μL of Opti-MEM. The diluted siRNA was then added to the diluted transfection reagent at a 1:1 ratio and allowed to incubate for 5 min at room temperature. 250 μL of the resulting siRNA-lipid complex was then added to each well for a total of 25 pmol siRNA + 7.5uL transfection reagent per well. Cells were incubated for 48 hr at 37 °C, 5% CO_2_. CAD5 cells were split at a 1:5 ratio 48 hr post-transfection and allowed to adhere for 24 hr complete media. After 24 hr, cells were subjected to re-transfection using the same methodology as above and were allowed to incubate for 72 hr at 37 °C, 5% CO_2_, after which cells were lysed. Western blot was performed on cell lysate to assess PrP^C^ as described above or to assess knockdown efficiency of *Gsk3a* or *Gsk3b* using anti-GSK-3α monoclonal antibody CST4337 (Cell Signaling Technology, Danvers, MA, USA) or anti-GSK-3β monoclonal antibody CST12456 antibody (Cell Signaling Technology) as primary antibodies and horseradish peroxidase-labeled goat-anti-rabbit secondary antibody (BioRad, Hercules, CA, USA, Cat # 1706515) at manufacturers’ recommended dilutions. The membrane was stripped, blocked, and relabeled using anti-beta Tubulin antibody CST15115 and horseradish peroxidase-labeled goat-anti-rabbit secondary antibody (BioRad Cat # 1706515).

### Generation of a *Gsk3b* knockout cells using CRISPR-Cas9

sgRNAs targeting *Gsk3b* were selected from the Brie library[15] and oligos (5’-CACCGACTGTAACATAGTCCGACTG-3 and 5’-AAACCAGTCGGACTATGTTACAGTC-3’) (Integrated DNA Technologies, Coralville, IA, USA) were used for cloning into the lentiGuide-Puro backbone as previously described[16] to generate specific lentiCRISPR plasmids, transformed into NEB Stable Competent *E. coli* (New England Biolabs, Ipswich, MA, USA), and verified using whole plasmid sequencing.

HEK293FT cells were used for lentiviral packaging of *Gsk3b* lentiCRISPR plasmids. HEK293FT cells were seeded 24 hr before transfection in 15 cm plates to allow for ∼60% confluence on the day of transfection. Medium containing G418 was removed 30 min before transfection and replaced with 25 mL of IMDM supplemented with 10% FBS, 6 mM L-glutamine, and 1X non-essential amino acids (transfection medium), with 12.5 μg lentiCRISPR DNA, 7.5 µg psPAX2, and 5 μg pMD2.G and gently vortexed. LipoD293 (SignaGen, Frederick, MD, USA) (75 μL) was added to 1.25 mL of the transfection medium and gently vortexed. This diluted LipoD293 was immediately added to the DNA mixture, gently vortexed, and incubated for 10 min at room temperature. The LipoD293/DNA mixture was added slowly to the cells with swirling for distribution, then incubated for 24 hr at 37 °C, 5% CO_2_. The medium was then replaced with IMDM (ThermoFisher) supplemented with 2% FBS (Cytiva), 6 mM L-glutamine (Gibco), 1X non-essential amino acids (Gibco), and 2 mM caffeine (MilliporeSigma) and incubated for an additional 24 hr. Lentiviral particles were harvested by filtration through a 0.45-μm filter and concentrated 10X using Amicon Ultra-15 10kDa Centrifugal Filter units (MilliporeSigma).

Monoclonal CAD5 cells constitutively expressing Cas9 were seeded in 150 mm plates 24 hr before transduction to allow for ∼60% confluence on the day of transduction. Cells were transduced with 250 μL lentivirus in the presence of 8 μg/mL polybrene. After 24 hr incubation, media was replaced with complete media. After 24 hr recovery, cells were kept under selection using 3.5 μg/mL puromycin in complete media. Monoclonal *Gsk3b* KO cells were isolated using serial dilution.

### Whole cell proteomics

Polyclonal *Gsk3b* KO CAD5 cells and WT CAD5 cells underwent mass spectroscopy using tandem mass tagging (TMT). Samples were run in biological triplicate. Cell pellets were lysed with urea and sonication and 30 μg of protein was removed for processing. After reduction, alkylation, and a methanol chloroform precipitation, extracted proteins were resuspended and digested with trypsin (1:100 w/w) overnight. Resulting peptides were desalted and labeled with TMT reagents (Thermo Scientific) prior to sample combination. Multiplexed samples were desalted and separated into 16 fractions by off-line HPLC prior to LCMS analysis. Peptides were separated across a 120-min gradient of 4-41% acetonitrile in 0.1% formic acid over a 30 cm x 100 µm C18 column (Dr. Maisch) with a Vanquish Neo (Thermo Scientific) liquid chromatography system and electrosprayed into an Orbitrap Fusion Lumos (Thermo Scientific) mass spectrometer. Precursor ion (MS1) scans were obtained at 120,000 resolution in centroid in the orbitrap. Fragment ion (MS2) scans were acquired in data dependent mode in the linear ion trap. Reporter ion scans (MS3) were obtained via synchronous precursor selection at 50,000 resolution in the orbitrap.

Raw data were searched against a target-decoy database containing the mus musculus proteome using Comet[17, 18] requiring tryptic cleavage, mass accuracy of ±5 ppm, 3 missed cleavages, and the following modifications: Met oxidation (variable), Cys carbamidomethylation (static), Lys and peptide N-terminus TMT labeling (static). Results were filtered to an FDR <1%. Peptide reporter ions were summed to obtain protein quantities. Proteins identified by fewer than 2 unique peptides were removed from the dataset. Resulting abundances were normalized to the total signal across all channels. Significant changes in protein abundance between conditions were assessed by two-sided moderated t-test using the limma package within the R framework[19]

### Treatment of CAD5 cells with pharmacological modulators of protein turnover

WT CAD5 or monoclonal *Gsk3b* KO cells were seeded in 6-well tissue culture plates (Corning) and allowed to grow to 60% confluence. Cells were treated with complete media containing either 1 μM MG-132 (MilliporeSigma Cat # M7449), 50 nM bafilomycin (MilliporeSigma Cat # SML1661), 10 μg/mL cycloheximide (MilliporeSigma Cat # C4859), or 10 μM laduviglusib for 24 hr.

## Acknowledgements

This study was funded by the National Institute for Neurological Disorders and Stroke (1R37NS125431, R01NS117276 and R01NS118796 to S.S.) and the National Institutes of Health (P30-GM165328 to Dean Madden). The authors acknowledge the Biological Mass Spectrometry and Proteomics Shared Resource at the Dartmouth Cancer Center (RRID:SCR_026076) with NCI Cancer Center Support Grant P30CA023108.

## Results

### *Gsk3b* is a positive regulator of PrP^C^ expression

To test the regulation of PrP^C^ levels by GSK paralogs, we first treated CAD5 cells with siRNA targeting *Gsk3a* and/or *Gsk3b* and used western blot to measure total PrP^C^ levels. The results show that *Gsk3a* knockdown alone has no effect on total PrP^C^ expression **(Figure 1A, middle panel, lanes 3-4)**, whereas *Gsk3b* knockdown reduces PrP^C^ levels either alone **(Figure 1A, middle panel, lanes 5-6)** or in combination with *Gsk3a* knockdown **(Figure 1A, middle panel, lanes 7-8)**. In addition, we generated monoclonal *Gsk3b* KO cells using CRISPR Cas9 and found that these KO cells had significantly lower levels of surface **(Figure 1B)** and total **(Figure 1C)** PrP^C^ than WT control cells. Taken together, these findings show that GSK-3β (but not its highly homologous paralog GSK-3α) is a positive regulator of both cell surface and total PrP^C^ levels in CAD5 cells.

**Figure 1:**
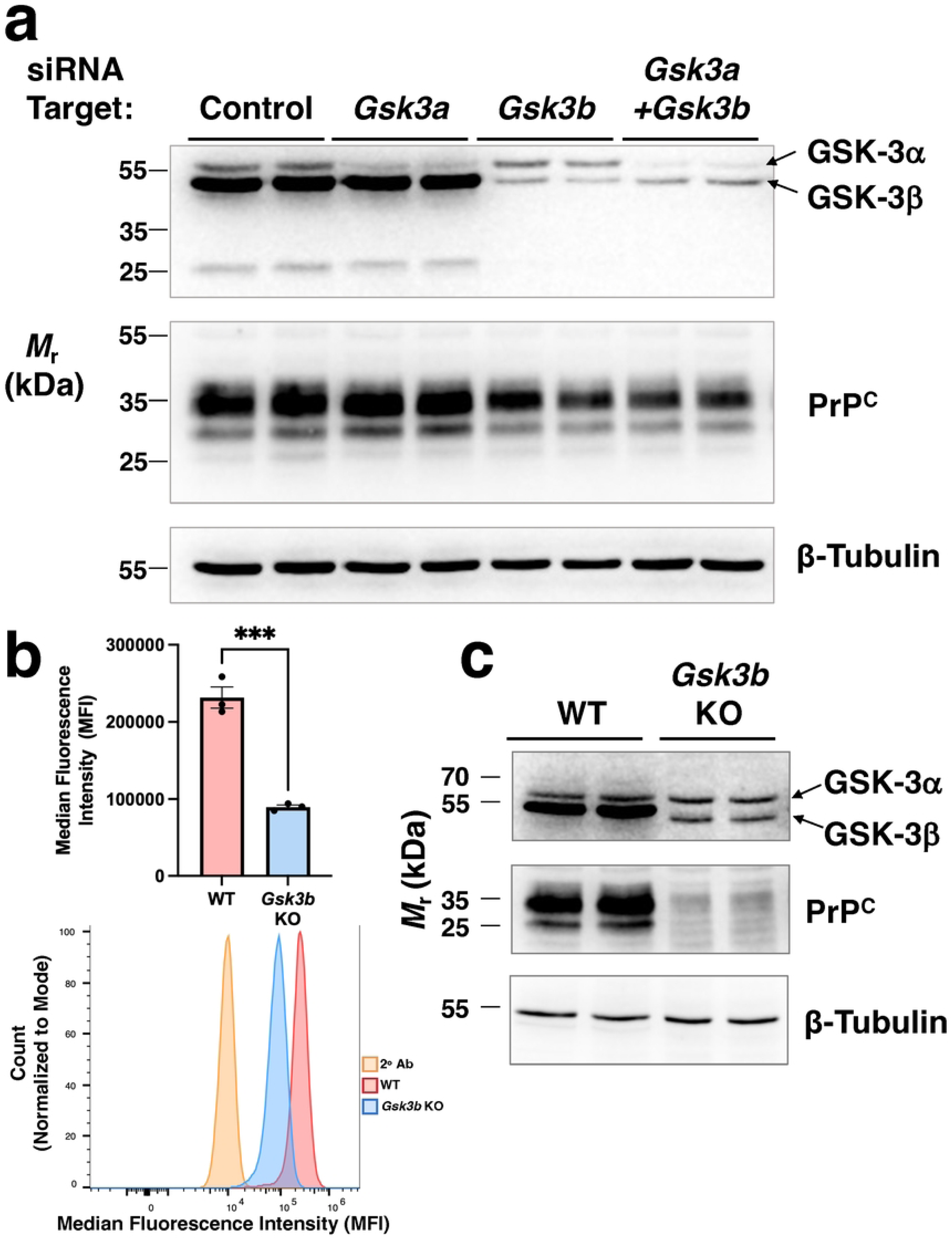
Genetic disruption of *Gsk3b* reduces PrP^C^ levels in CAD5 cells. **(A)** Western blot of crude lysates of cells treated with siRNA targeting *Gsk3a* and/or *Gsk3b*, as indicated. A single blot was sequentially probed with antibodies against GSK-3, PrP, and β-tubulin, as indicated. Relative migration of GSK-3 paralogs indicated with arrowheads. **(B)** Quantification (Top panel) and representative histograms (Bottom panel) of cell surface PrP^C^ levels in WT and monoclonal *Gsk3b* knockout (KO) cells, measured by flow cytometry following staining with anti-PrP mAb 6D11 (n = 3 biological replicates, *** p-value < 0.001). **(C)** Western blot of crude lysates of WT and *Gsk3b* KO cells, as indicated.

### PrP^C^ is one of the most significantly downregulated proteins in GSK-3β knockout cells

To visualize at a global level how GSK-3β impacts PrP^C^ levels relative to the levels of other cellular proteins, we performed whole cell proteomics on *Gsk3b* polyclonal KO (a population containing a mixture of complete and partial CRISPR knockouts) and WT CAD5 cells and quantitatively compared protein abundances using tandem mass tagging (TMT) **(Figure 2)**. Strikingly, the results show that PrP^C^ is one of the most significantly downregulated proteins in *Gsk3b* KO cells relative to WT cells (rank #60 out of 7227 detected proteins, p < 0.000013) **(Figure 2, red circle labeled “PrP^C^”)**.

**Figure 2:**
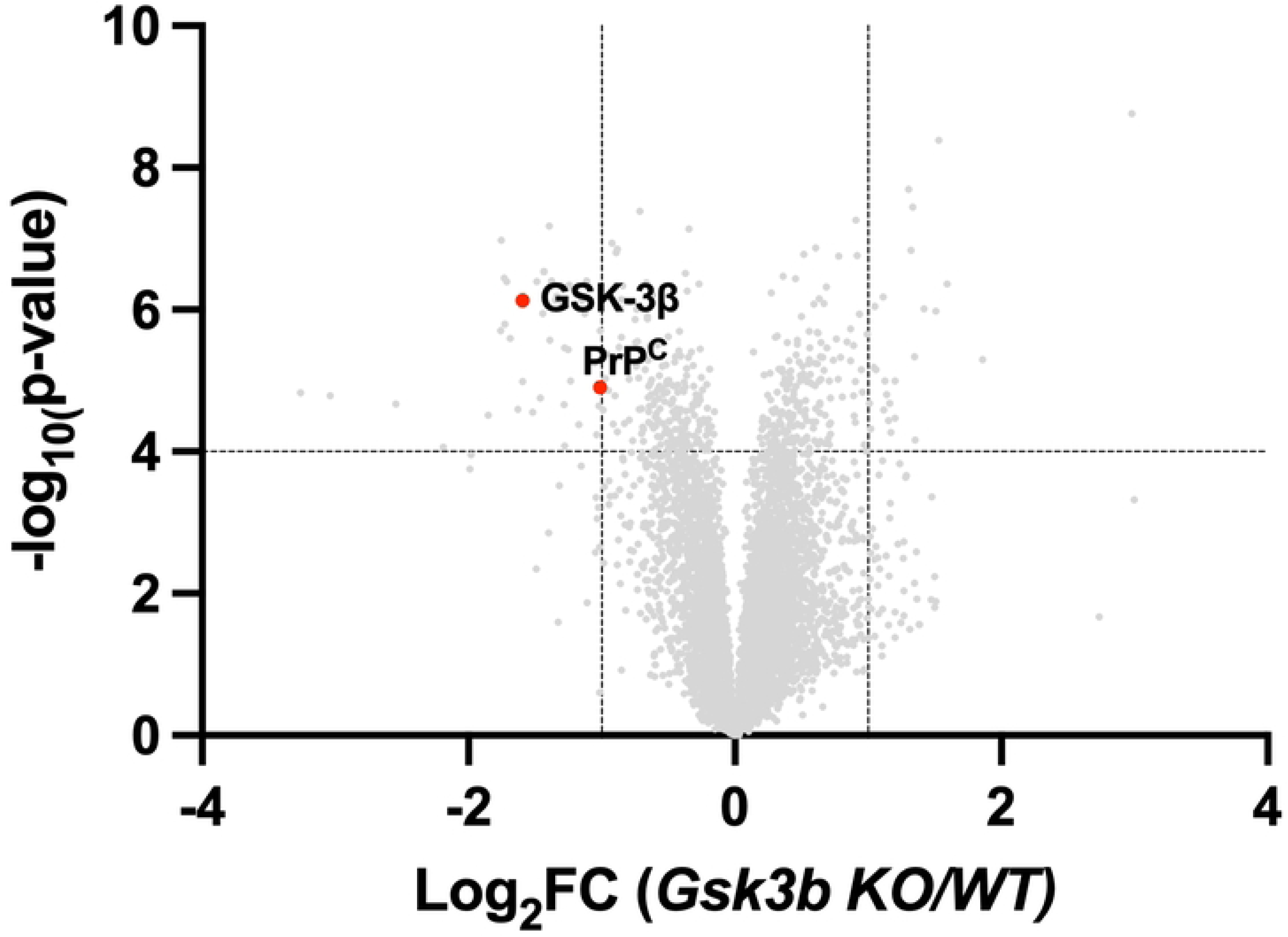
Whole cell proteomic analysis of *Gsk3b* knockout cells. Volcano plot showing changes in protein abundance between polyclonal *Gsk3b* knockout and WT CAD5 cells. This experiment was performed in biological triplicate.

### Small molecule GSK-3 inhibitors reduce PrP^C^ levels in undifferentiated and differentiated mouse and human cells

Since GSK-3β is an attractive drug target for AD and a variety of other neurological conditions, several pharmaceutical companies have developed orally available, brain-penetrant small molecule GSK-3 inhibitors [10, 11]. Therefore, we evaluated two commercially available compounds, laduviglusib (CHIR-99021) and AZD-1080, for their ability to modulate cell surface PrP^C^ levels, as measured by flow cytometry. We tested these compounds in CAD5 cells and human BE(2)-C neuroblastoma cells, both in their undifferentiated and differentiated states **(Supplemental Figure S1)**. In both states of differentiation for each cell line, both compounds effectively reduced surface PrP^C^ levels **(Figure 3A)**. Total PrP^C^ levels were also reduced by drug treatment **(Figure 3B)**, and the magnitude of the drug-induced PrP^C^-lowering effect was dependent on both the dose and duration of treatment **(Supplemental Figure S2)**.

**Figure 3:**
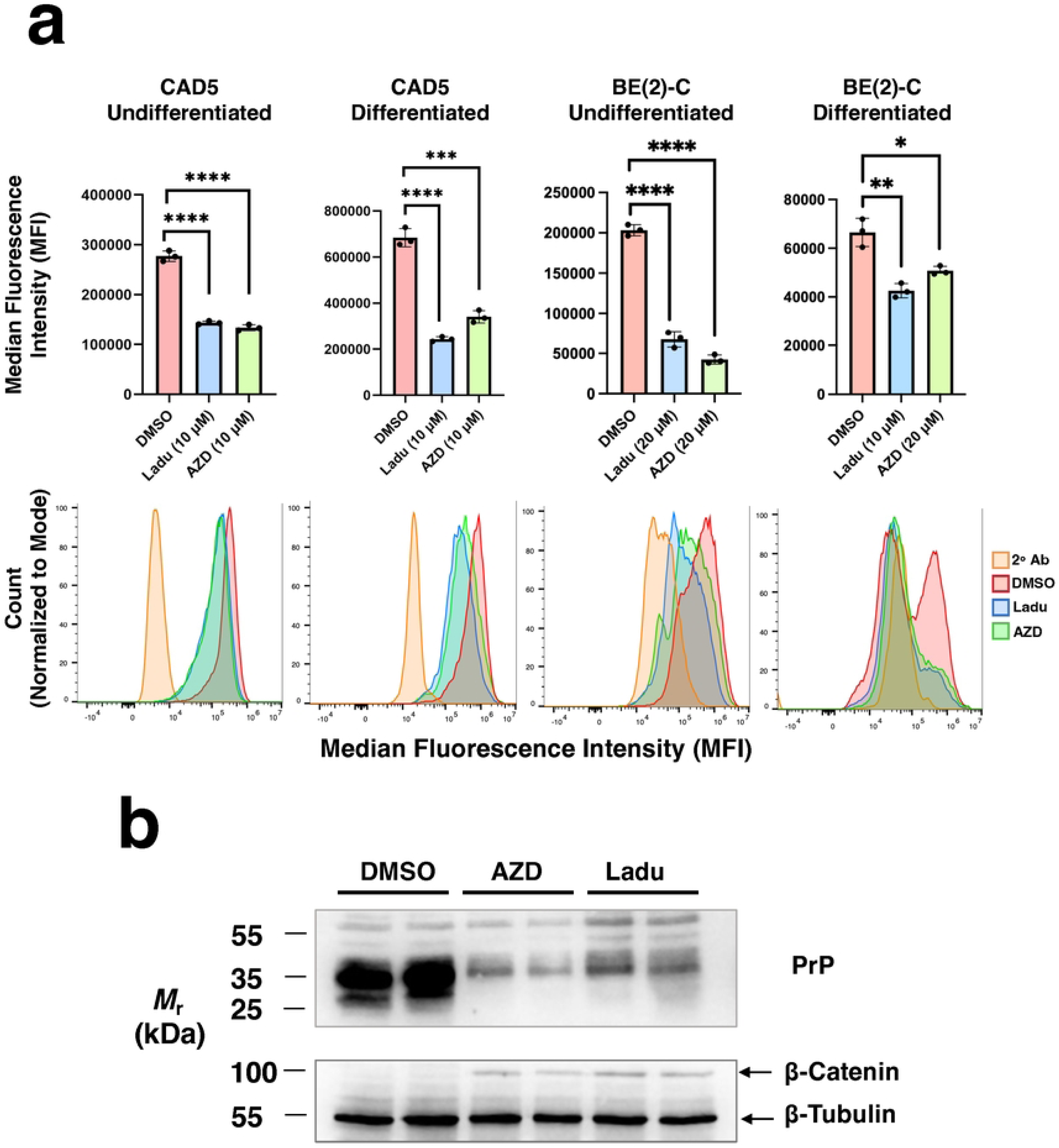
Effect of small molecule GSK-3 inhibitors on surface PrP^C^ levels. **(A)** Quantification (Top) and representative graphs (Bottom) of cell surface PrP^C^ levels in undifferentiated and differentiated CAD5 and BE(2)-C cells, measured by flow cytometry following staining with anti-PrP mAb 6D11 (n = 3 biological replicates, * p-value < 0.05, ** p-value < 0.01, *** p-value < 0.001, **** p-value < 0.0001). Cells were treated for 48 hr, as indicated. Ladu = cells treated with laduviglusib, AZD = cells treated with AZD-1080, 2° Ab = secondary antibody only staining control. **(B)** Western blot of crude lysates of undifferentiated CAD5 cells treated with DMSO, 10 µM AZD-1080, or 10 µM laduviglusib for 48 hr, as indicated. A single blot was sequentially probed with antibodies against PrP and β-tubulin, as indicated.

### GSK-3β regulates PrP^C^ levels through a post-transcriptional process

We performed two sequential experiments to investigate the mechanism by which GSK-3β inhibition reduces PrP^C^ levels. First, we tested whether GSK-3β inhibition could reduce surface PrP^C^ levels in CAD5-PrP^-/-^(BV^M^) cells, which lack endogenous mouse PrP^C^ and instead express bank vole PrP^C^ from a CMV promoter[20]. The results show that laduviglusib successfully reduced surface PrP^C^ in CAD5-PrP^-/-^(BV^M^) cells, as judged by flow cytometry **(Figure 4A-B)**, suggesting that PrP^C^ levels are reduced via a mechanism that does not involve transcriptional regulation of the *Prnp* promoter. Next, we treated WT and *Gsk3b* KO CAD5 cells with pharmacological modulators of protein turnover to identify the post-transcriptional processes that mediate GSK-3β regulation of surface PrP^C^ levels, as determined by flow cytometry. The results show that MG132, a proteasome inhibitor, increased surface PrP^C^ levels in *Gsk3b* KO cells **(Figure 4D, compare black bar to white bar)**, but not in WT cells **(Figure 4C, compare black bar to white bar)**. Bafilomycin (a lysosome and autophagy inhibitor) had no effect on PrP^C^ levels in *Gsk3b* KO cells **(Figure 4D, compare dark grey bar to white bar)**, but surprisingly reduced PrP^C^ levels in WT cells **(Figure 4C, compare dark grey bar to white bar)**. Cycloheximide (a protein translation inhibitor) reduced PrP^C^ levels by a similar percentage in *Gsk3b* KO (DMSO/CHX = 54.5%) and WT cells (DMSO/CHX = 59.2%) **(Figures 4C-D, compare light grey bars to white bars)**, indicating that GSK-3β impairs PrP^C^ turnover post-translationally. Taken together, these results indicate that GSK-3β regulation of surface PrP^C^ levels occurs post-translationally, possibly involving both proteasomal and lysosomal degradation.

**Figure 4:**
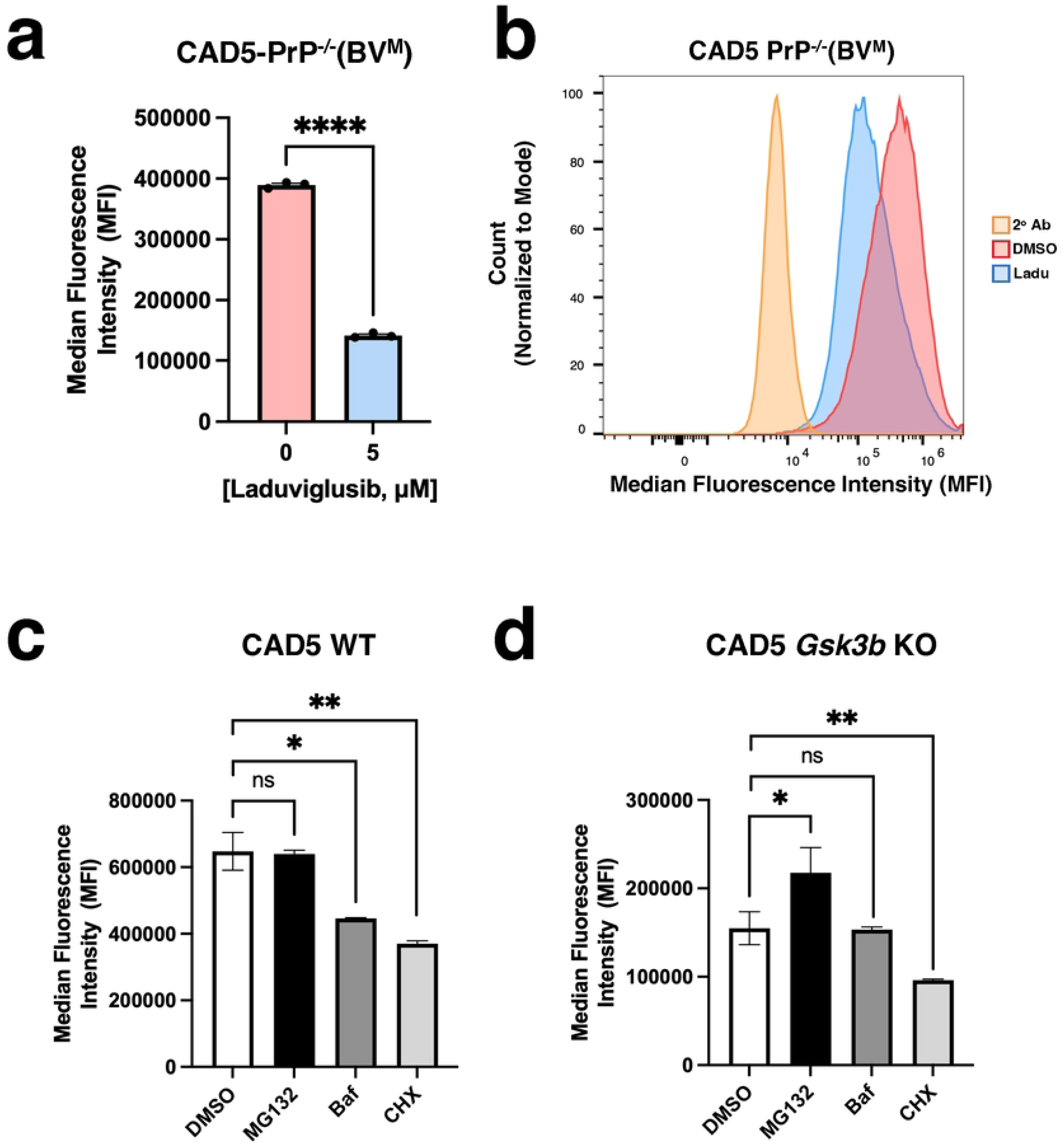
GSK-3β regulates PrP^C^ levels through a post-translational mechanism. **(A)** Quantification and **(B)** representative flow cytometry histograms of surface PrP^C^ levels in CAD5-PrP^-/-^(BV^M^) cells treated with either laduviglusib (Ladu) or AZD-1080 (AZD), as indicated (n = 3 biological replicates, **** p-value < 0.0001). **(C-D)** Quantification of surface PrP^C^ levels (determined by flow cytometry following staining with anti-PrP mAb 6H11) of **(C)** WT and **(D)** *Gsk3b* KO cells treated with DMSO, MG132, bafilomycin (Baf), or cycloheximide (CHX) for 24 hr, as indicated. (n = 3 biological triplicates, *p-value < 0.05, ** p-value < 0.01 unpaired t-test. ns = not significant).

### Small molecule GSK-3 inhibitors reduce PrP^Sc^ in CAD5 cells chronically infected with hamster and mouse prions

Since PrP^C^ is an obligate substrate for the production of all PrP^Sc^ conformers, we reasoned that small molecule GSK-3 inhibitors might reduce PrP^Sc^ levels in cells infected with various prion strains. To test this possibility, we treated prion-infected CAD5 cells with either laduviglusib **(Figure 5A, top blot)** or AZD-1080 **(Figure 5A, bottom blot)** and measured PrP^Sc^ levels in crude cell lysates by proteinase K digestion and western blot. The results show that both drugs successfully reduced PrP^Sc^ levels in CAD5 cells chronically infected with three different prion strains: Hyper hamster prions **(Figure 5A-B, lanes 1-4)**, 22L mouse prions **(Figure 5A, lanes 5-8)**, and RML mouse prions **(Figure 5A, lanes 9-12)**.

**Figure 5:**
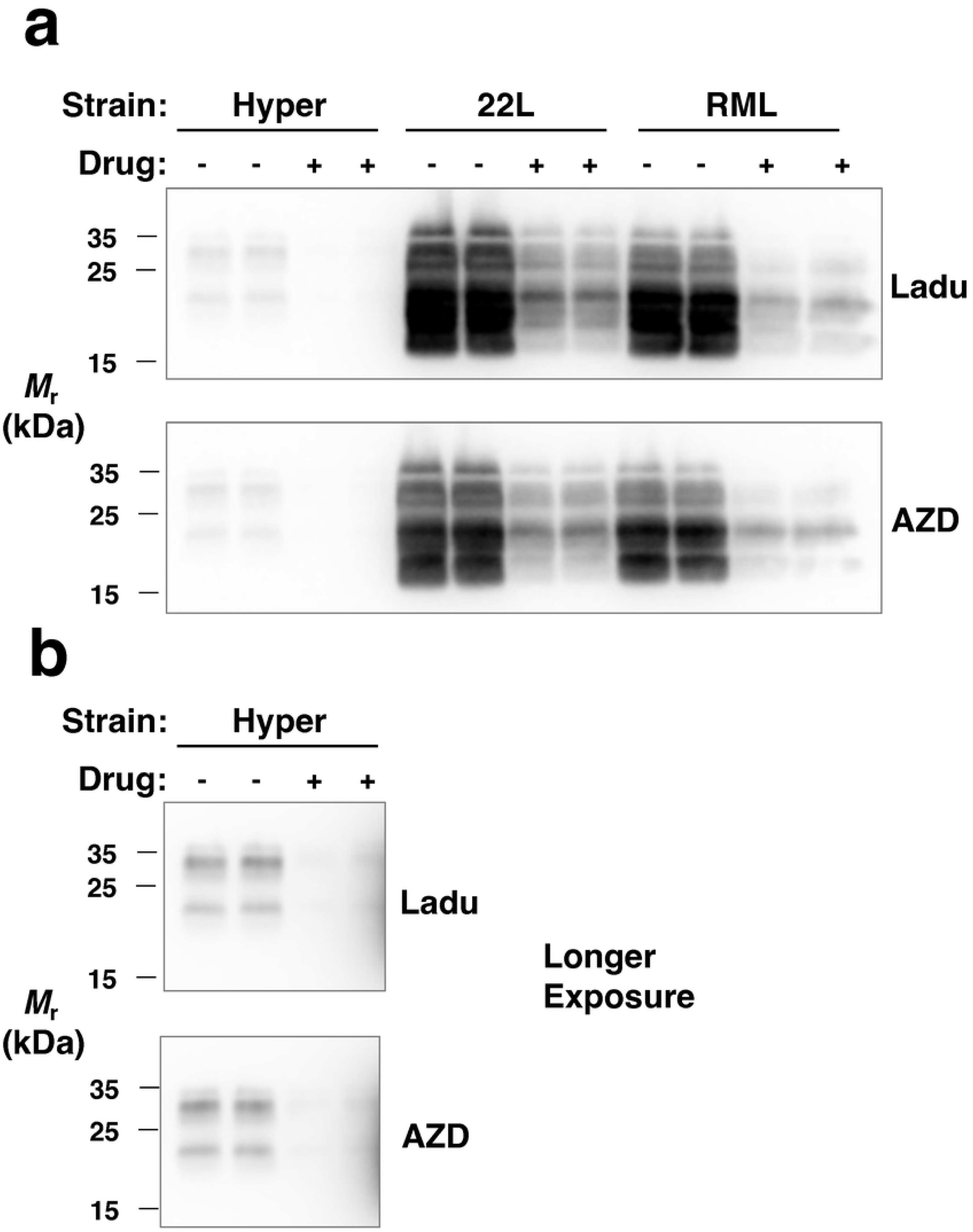
Small molecule GSK-3 inhibitors reduce PrP^Sc^ levels in CAD5 cells infected with various prion strains. **(A)** Western blots of crude lysates of CAD5-PrP^-/-^(Ha) cells infected with Hyper hamster prions and WT CAD5 cells infected with either 22L or RML mouse prions, as indicated. Cells were treated with DMSO, 10 µM laduviglusib (Ladu, top blot), or 10 µM AZD-1080 (AZD, bottom blot) for 72 hr, as indicated. All samples were treated with proteinase K and the blot was probed with anti-PrP mAb 6D11. **(B)** Longer exposure to show faint bands in Hyper-infected CAD5-PrP^-/-^(Ha) cells more clearly.

## Discussion

In this study, we report that GSK-3β is a positive regulator of cellular PrP^C^ levels. Either genetic disruption of *Gsk3b* expression or pharmacological inhibition of GSK-3β activity decreased PrP^C^ levels in both mouse and human cells. These findings reveal a novel, post-translational mechanism that controls PrP^C^ levels.

### PrP^C^ is a logical therapeutic target for prion diseases

It is known that reducing PrP^C^ levels is an effective strategy for treating prion diseases. Knockout mice lacking PrP^C^ are resistant to prion infection and reducing neuronal PrP^C^ levels extends the survival of prion-infected animals[21, 22]. For instance, lowering PrP^C^ by 50% increases the survival time in scrapie-infected mice ∼4-fold[23, 24]. In addition to being a required substrate for prion disease, PrP^C^ is also a high-affinity receptor for Aβ oligomers[2]. Binding of Aβ oligomers to PrP^C^ activates Fyn kinase and impairs synaptic function in AD mouse models[4, 25, 26].

Targeting PrP^C^ levels circumvents several problems associated with anti-PrP^Sc^ drugs, e.g. prion strain-specificity, drug resistance, and ineffectiveness against inherited forms of prion disease[27–31]. In this study, we show that two different GSK-3β inhibitors can reduce PrP^Sc^ levels in cells infected with three different prion strains. These results suggest that GSK-3β inhibition might be a versatile strategy that could be used to treat various types of prion disease.

### GSK-3β modulates PrP^C^ levels through a post-translational mechanism

Cells maintain PrP^C^ levels through an intricate set of processes including transcription, translation, post-translational modifications, membrane trafficking, and enzymatic cleavage. Transcription of the *Prnp* gene is mediated by a TATA-less promoter (containing Sp1 sites) as well as non-coding regions, and can be influenced by factors such as cell stress[32, 33] and ERK1 activity[34]. Following translation, PrP^C^ is translocated into the endoplasmic reticulum through the Sec62/63 complex and subsequently trafficked to cell surface lipid rafts (a process that requires addition of a glycophosphatidylinositol (GPI) anchor)[8, 35]. Cell surface PrP^C^ levels are also influenced by N-glycosylation of PrP residues Asn181 and Asn197[36]. From the cell surface, PrP^C^ can be endocytosed, recycled[37], or shed by ADAM17 metalloprotease [34].

Our results suggest that GSK-3β regulates PrP^C^ levels through a post-translational mechanism, since cycloheximide reduced PrP^C^ levels similarly in *Gsk3b* KO and WT cells. Furthermore, PrP^C^ levels in *Gsk3b* KO cells were partially rescued by MG132 and maintained by bafilomycin, suggesting that GSK-3β might modulate the level PrP^C^ clearance by both proteasomes and lysosomes.

### GSK-3β is a paralog-specific and potent regulator of PrP^C^ levels

GSK-3 is a highly conserved serine/threonine kinase known to phosphorylate many substrates, including tau, β-catenin, and CREB, linking it to the pathogenesis of neurodegenerative diseases, cancer, and psychiatric disorders[9]. Originally identified for its role in glycogen metabolism, GSK-3 has subsequently been implicated in diverse physiological processes including Wnt/β-catenin signaling, insulin signaling, circadian rhythm regulation, neurodevelopment, and apoptosis[38, 39]. There are two paralogs of GSK-3 (GSK-3α and GSK-3β) encoded by distinct genes but sharing significant structural and functional similarity, particularly within their kinase domains[40]. Despite these similarities, the two paralogs do exhibit some differences in substrate specificity[41]. Our siRNA and CRISPR KO data show that GSK-3β is the only paralog responsible for regulating PrP^C^ levels. This specificity could potentially be exploited therapeutically through the use of inhibitors that block GSK-3β but not GSK-3α. Strikingly, our whole cell proteomics data indicate that PrP^C^ is one of the most significantly downregulated proteins in *Gsk3b* KO versus WT cells. Taken together, the paralog specificity and magnitude of its effect on PrP^C^ suggest that GSK-3β is likely to be a physiological regulator of PrP^C^ levels.

### GSK-3β is a potential drug target

Currently, there are no FDA-approved treatments available for patients with prion disease. Attempts to identify small molecules that inhibit the conversion of PrP^C^ to PrP^Sc^ through high-throughput screening have been plagued by species-and strain-specificity as well as drug resistance[27, 28, 42]. Furthermore, drugs that inhibit WT PrP^Sc^ formation appear to have no effect in mouse models of inherited prion disease[30]. Alternative therapeutic strategies designed to reduce or block PrP^C^ expression also have specific logistical drawbacks. Anti-PrP^C^ antibodies have poor brain penetration and may have toxic side effects on neurons[43, 44]. Anti-sense oligonucleotides (ASOs) must be repeatedly administered by lumbar puncture and show varying degrees of effectiveness, especially in deep brain regions such as the thalamus and hippocampus[6]. Gene therapies that depend on viral packaging may not be successfully delivered to all the neurons in the brain and could potentially induce irreversible off-target effects[45].

An ideal drug for prion diseases would be an orally available, brain-penetrant, well-tolerated small molecule that lowers surface PrP^C^ levels in neurons. To our knowledge, no such drug has been identified to date. A prior screen of 1040 repurposed drugs identified FK506 as an oral compound that could reduce cellular PrP^C^ levels, but only at toxic concentrations[46]. A more recent screen of ∼3500 small molecules identified 2 compounds (EYH and LCZ) that successfully lowered PrP^C^ levels in mouse N2a cells, but unfortunately neither compound penetrated the BBB nor was effective in human U251-MG cells[47]. In contrast, our data show that GSK-3β inhibitors lower PrP^C^ levels in both mouse and human cells and in both undifferentiated and differentiated states. Furthermore, GSK-3β inhibitors could potentially be combined with other PrP^C^-lowering therapies, such as ASOs[48] to simultaneously block PrP^C^ biosynthesis and increase PrP^C^ degradation.

Due to its ability to promote tau hyperphosphorylation, Aβ production, mitochondrial fragmentation, and neuroinflammation while simultaneously suppressing neurogenesis and neuroplasticity; GSK-3β has emerged as a promising therapeutic target, particularly for CNS disorders such as AD[49, 50]. As a result, many orally bioavailable, brain-penetrant small-molecule GSK-3 inhibitors have been developed. Many inhibitors have caused unacceptable side effects such as hypoglycemia, due to the presence of pharmacological targets other than PrP^C^[10–12]. Nonetheless, some compounds, such as laduviglusib, Elraglusib (9-ING-41), and tideglusib, have been relatively well-tolerated and advanced to clinical trials (NCT numbers: 03678883, 05086276, 05004129), and many new inhibitors are being actively developed, including highly-specific substrate-competitive inhibitors[51].

## Limitations

Although this study establishes GSK-3β as an potent regulator of PrP^C^ levels, it is important note several experimental limitations. (1) We have shown that GSK-3β regulates PrP^C^ levels in both mouse and human cells, but additional work is required to test its importance in other species and in whole organisms. (2) All the experiments conducted in this study were relatively short term, ranging from 1-3 days. Additional experiments are needed to determine whether PrP^C^ levels remain depressed after longer periods of GSK-3β inhibition, or whether compensatory mechanisms might eventually restore PrP^C^ levels. (3) We have shown that GSK-3β regulates PrP^C^ levels through a post-translational mechanism that may involve proteasomes, but have not yet elucidated the molecular details of this process. (4) Finally, we have shown that PrP^C^ regulation is paralog-specific (i.e. only GSK-3β but not GSK-3α influences PrP^C^ levels), but we have not yet identified the GSK-3β-specific phosphosubstrate(s) responsible for this effect.

## Conclusion

Through a combination of genetic and pharmacological experiments, we show for the first time that GSK-3β is a positive regulator of PrP^C^ levels in both mouse and human cells, in both undifferentiated and differentiated states. This discovery reveals a novel post-translational process for regulating PrP^C^ levels through kinase signaling with translational potential.

## Supplemental Figure Legends

**Figure S1.** (B) Representative images of undifferentiated (left) and differentiated (right) CAD5 (top) and BE(2)-C (bottom) cells.

**Figure S2.** Graphs of cell surface PrP^C^ levels in CAD5 cells (as detected by flow cytometry in biological triplicate) following treatment with laduviglusib at various concentrations and incubation times, as indicated.

